# The blue dragon nudibranch (*Glaucus atlanticus*) shifts to less venomous prey in the Mediterranean, with no evidence of defensive microbial symbionts

**DOI:** 10.64898/2026.11.24.768820

**Authors:** Dafne Porcel Sanchis, Alicia Ortiz-Maiques, Inés Alberola Mora, María Asunción Piedecausa Narejos, Conxita Avila, Itziar Álvarez-Trasobares, Laura Prieto, Mária Džunková

## Abstract

*Glaucus atlanticus* (“blue dragon”), is an aeolid nudibranch that acquires chemical defense through sequestration of nematocysts from its primary prey, the highly venomous *Physalia physalis* (“Portuguese man-of-war”). In August 2025 hundreds of *G. atlanticus* individuals appeared near Alicante (Spain), prompting temporary beach closures due to concerns about potential envenomation. This unusual occurrence, in a region where *P. physalis* is typically absent, raises questions about the trophic ecology and defensive strategies of this species under novel environmental conditions.

Here, we used COI metabarcoding of gut to characterize the diet of *G. atlanticus* and combined it with 16S rRNA gene sequencing of cerata, gut, mucus, and central dorsal epidermis to assess potential microbial transfer from alternative prey that could contribute to the synthesis of defensive compounds, given that the role of microbes in chemical defense of aeolid nudibranchs was not assessed previously.

We detected a trophic shift toward alternative cnidarian prey, including *Pelagia noctiluca* and *Porpita porpita*, which are not considered highly hazardous. Microbiome comparisons with *P. porpita*, as well as comparison of core microbiomes across different body parts of *G. atlanticus*, revealed no evidence of prey-derived transfer of mutualistic bacteria. Instead, *G. atlanticus* harbored distinct microbial communities dominated by environmentally widespread taxa, including potential opportunistic pathogens, with no indication of symbionts associated with defensive functions.

Altogether, these findings indicate that, despite trophic flexibility, *G. atlanticus* may have limited capacity to maintain its defensive mechanisms in the absence of its primary prey.

## Introduction

*Glaucus atlanticus* Forster, 1777 (“blue dragon”) is a holoplanktonic nudibranch mollusc (Thompson and Bennett, 1969; Valdes and Angulo, 2004), widely distributed in tropical and subtropical oceans worldwide, where it inhabits the ocean surface and drifts with pleustonic communities (Pinotti et al., 2019). In August 2025, hundreds of individuals of *G. atlanticus* appeared along the Spanish Mediterranean coast, particularly in Guardamar del Segura (Alicante, Valencia) (**Figure 1a**), prompting the temporary closure of local beaches attracting considerable media attention (Artigas, 2025; Sánchez, 2025; Wolfe, 2025). This precautionary measure was related to the potential danger posed by this species, which derives from its most common prey, the hydrozoan *Physalia physalis* Linnaeus, 1758 (“Portuguese man-of-war”), whose sting can cause severe and, in rare cases, potentially life-threatening envenenomation in humans (Labadie et al., 2012). *P. physalis* is highly venomous due to the presence of nematocysts, specialized organelles in cnidarians capable of discharging bioactive compounds stored within the capsule matrix, which is used for prey capture and defense (Özbek, 2011). Those nematocysts can be sequestered and stored within specialized nudibranch cells (cnidosacs) in the cerata of *G. atlanticus*. of *G. atlanticus*. Cerata are dorsal appendages containing gastrodermal channels of the digestive gland that also serve as storage of prey-derived defensive structures (Vorobyeva et al., 2023). The sequestered nematocysts, known as kleptocnides (stolen nematocysts), allow the nudibranch to retain functional stinging cells (Edmonds, 1995; Johnson and Allen, 2005). As a result, *G. atlanticus* may serve as a vector for *P. physalis* venom (Yamamoto et al., 2025), representing a hypothetical public health risk.

**Figure 1.**
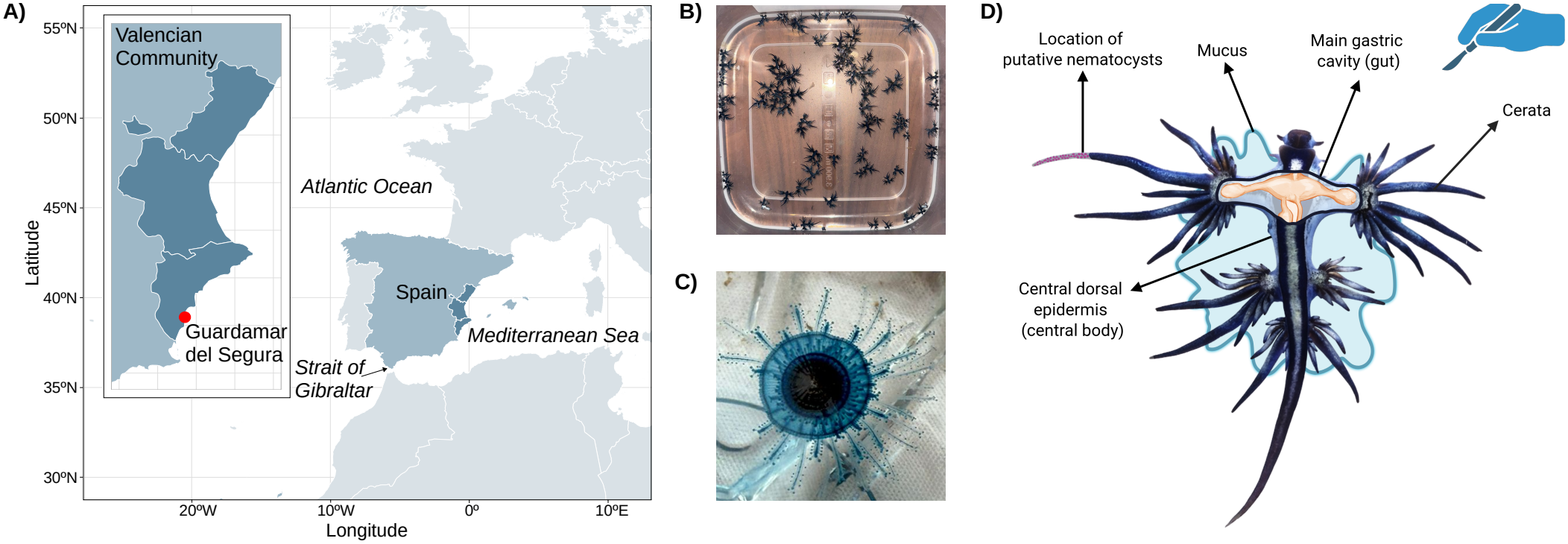
Study design. **a)** Sampling location in Guardamar del Segura. **b)** Example of the number of *G. atlanticus* individuals collected daily by beach guards in August 2025. **c)** *P. porpita*, Guardamar del Segura. **d)** Schematic representation of *G. atlanticus* dissection and sampled body parts.

Although *G. atlanticus* was first reported in the Mediterranean in the 18th century (Forster, 1777), no further reports were documented for more than 300 years, until it was sporadically observed again in 2021 and 2024 (Chebaane et al., 2024; Pujol et al., 2023). The mass occurrence (**Figure 1b**) of this generally uncommon nudibranch along the Spanish coasts in August 2025 was likely facilitated by Atlantic currents and increasingly warmer Mediterranean waters (Gonzalez et al., 2025; Schroeder et al., 2016). Its most common prey, *P. physalis*, can be transported by the same currents (Chebaane et al., 2024; Macias et al., 2025; Tiralongo et al., 2022); however, its occurrence near the same beaches in the Alicante Province has not been documented. Beyond *P. physalis, G. atlanticus* has been reported to feed on the hydrozoans *Velella velella* (“by-the-wind sailor”) and *Porpita porpita* (“blue button”, **Figure 1c**), and occasionally fish (Helm, 2021; Yamamoto et al., 2025). These prey species, unlike *P. physalis*, are commonly observed near Alicante beaches and do not lead to beach closures since they are less dangerous to humans. We therefore hypothesize that the *G. atlanticus* individuals found at these locations do not feed on *P. physalis*, but instead consume these less common food sources, which may result in the sequestration of different compounds that are potentially less harmful to humans.

Nevertheless, the investigation of defensive compounds acquisition in nudibranchs cannot be limited to the transfer of nematocysts alone, as some species rely on microbial symbionts for the production of defensive compounds (Gris and Prinsep, 2026). For example, the dorid nudibranch *Doriopsilla fulva* produces chemical defense molecules through its symbiont *Candidatus Doriopsillibacter californiensis* (Džunková et al., 2023). To date, no microbial symbionts have been firmly established for aeolid nudibranchs, such as *G. atlanticus*, which, in contrast to dorids, are known to sequester nematocysts (Ekimova et al., 2022; Goodheart et al., 2025, 2022). However, this apparent absence of mutualistic microbes may reflect the limited number of microbiome studies conducted. To our knowledge, only *Berghia stephanieae* (Sickinger et al., 2024) and *Pteraeolidia semperi* (Ng et al., 2022) have been investigated, with microbiomes dominated by *Mycoplasma*, a taxon associated with stress conditions (Sickinger et al., 2024), providing no clear evidence for symbionts involved in chemical defense. Regarding potential microbial transfer from the prey, in *B. stephanieae* the prey was examined and found to lack *Mycoplasma*, whereas in *P. semperi* the prey was not assessed, leaving the possibility of microbial transfer unresolved. Therefore, the recent mass occurrence of *G. atlanticus* in the Mediterranean provides a unique opportunity to investigate potential microbiome transfer from novel prey in aeolid nudibranchs.

Here, we combine COI metabarcoding and 16S rRNA gene sequencing across multiple body regions to characterize the diet of *G. atlanticus* and determine whether trophic shifts in the Mediterranean are accompanied by the acquisition of prey-derived microbes that could contribute to chemical defense. We analyzed microbiomes of different body parts of nine *G. atlanticus* individuals to identify consistently associated bacterial taxa (core microbiome), which is a necessary first step towards assessing a potential microbial contribution to chemical defence. Under the hypothesis that defensive compounds could be produced by bacterial symbionts, taxa showing high abundance and prevalence across *G. atlanticus* individuals in their cerata, mucus or even digestive tract may represent candidate symbionts warranting further investigation. This work constitutes the first comprehensive exploration of food sources and microbiome of an aeolid nudibranch, providing a basis for understanding its defensive compounds acquisition strategies.

## Methods

### DNA extraction

Nine specimens of *G. atlanticus* and three specimens of its putative prey *P. porpita* were collected from multiple locations along Centro Beach, Guardamar del Segura (Alicante, Spain, N 38° 4’ 51.068’’, W 0° 38’ 50.431’’), in August 2025 (**Supplementary Table S1**). Fresh specimens were immediately frozen at -20ºC after collection. Tissue samples from *P. porpita* and the central dorsal epidermis (central body), main gastric cavity (gut), and cerata of *G. atlanticus* were obtained by dissecting fragments (**Figure 1d**) of approximately 2 mm x 4 mm x 1 mm and subsequently homogenized in a Potter-Elvehjem tissue homogenizer (Fisher Scientific, Ref: 15301331) in 1 mL of saline solution (0.9% NaCl). The homogenate was split into halves. The first half of the resulting homogenates, together with the mucus samples, were filtered using a 5 µm filter to remove host-derived cells and analyze the bacterial fraction. The second half was left unfiltered for the analysis of eukaryotic cells. Genomic DNA was extracted using an alkaline lysis protocol used in previous studies (Porcel Sanchis et al., 2026). For 200 µL cell suspensions, 140 µL of lysis buffer (700 µL KOH [0.43 g/10 mL], 430 µL DTT [0.8 g/10 mL], and 340 µL nuclease-free water; pH 12) was added, and samples were incubated at room temperature for 10 min. Samples were then subjected to a thermal shock by freezing at −80 °C and thawing at 55 °C to promote lysis. The reaction was neutralized with 140 µL Tris-HCl buffer (0.5 g/mL, pH 4). DNA was subsequently treated with 1 µL proteinase K (20 mg/mL) at 55 °C for 30 min, purified using magnetic bead cleanup (Labclinics, Ref. CNGS-0050), and resuspended in 20 µL nuclease-free water.

### COI gene amplicon sequencing

The cytochrome c oxidase subunit I (COI) gene (Hebert et al., 2003) was amplified in 25 µL reaction volume using KAPA2G Fast HotStart ReadyMix (Merck, ref. KK5601), 1 µl of extracted DNA, and 5 µl of 1µM universal primers for marine invertebrates (Heller et al., 2018) containing overhang for Illumina library preparations: 5’-TCG TCG GCA GCG TCA GAT GTG TAT AAG AGA CAG GGW ACW GGW TGA ACW GTW TAY CCY CC-3’ and 5’-GTC TCG TGG GCT CGG AGA TGT GTA TAA GAG ACA GTA IAC YTC IGG RTG ICC RAA RAA YCA-3’. The following PCR conditions were used: initial denaturalization 95 °C for 3 min, followed by 33 cycles of 95 °C for 30 s, 48 °C for 30 s, and 72 °C for 30 s, final elongation at 72 °C for 5 min. The purified amplicons were sequenced on the Illumina MiSeq platform using a paired-end 2 × 250 bp protocol. In order to decrease the proportion of host DNA in the resulting amplicon datasets, sequencing was repeated using the same DNA extracts with a modification of the PCR reaction implementing *G. atlanticus* blocking primers. The blocking primers were designed using Primedigital (Kalendar, 2025) to match the melting temperature of the regular primers used in the same PCR reaction. The resulting sequences 5’ GGT ACT GGT TGA ACT GTT TAT CCT CC - C3spacer 3’ and 5’ TAG ACT TCG GGG TGG CCG AAG AAT CA -C3spacer 3’ were checked by aligning them to reads from the first sequencing run to confirm that the blocking primers did not interfere with prey detection. Ratio of 5:1 of blocking to universal primers was chosen, as it represented the minimum proportion at which the blocking primers showed effective performance (**Supplementary Figure S1**). PCR conditions used were the same as those for universal COI PCR described previously.

To demonstrate that our COI sequencing approach is suitable for detecting *P. physalis* DNA in *G. atlanticus* samples, we prepared positive controls by extracting DNA from P. physalis collected at Camposoto Bach, San Fernando (Cádiz, Spain, N 36° 25’ 35.890’’, W 6° 13’ 49.195’’) on the Spanish Atlantic coast and mixing it with DNA extracted from *G. atlanticus* specimens. For each PCR reaction, 1 ng of *G. atlanticus* DNA was mixed with *P. physalis* DNA at 100-, 1,000-, 10,000-, 100,000-, and 1,000,000-fold dilutions. The resulting mixtures were amplified and sequenced on the Illumina MiSeq platform as described above.

### 16S rRNA gene amplicon sequencing

The 16S rRNA gene was amplified using the same general approach as described above for the COI gene, with universal bacterial primers containing Illumina overhangs: 341F (5’-TCG TCG GCA GCG TCA GAT GTG TAT AAG AGA CAG CCT ACG GGN GGC WGC AG-3’) and 785R (5’-GTC TCG TGG GCT CGG AGA TGT GTA TAA GAG ACA GGA CTA CHV GGG TAT CTA ATC C-3’) (Klindworth et al., 2013). PCR conditions were as follows: initial denaturation at 95 °C for 3 min; 30 cycles of 95 °C for 30 s, 55 °C for 30 s, and 72 °C for 30 s; and a final elongation at 72 °C for 5 min. Purified 16S amplicons were sequenced on the same Illumina MiSeq run together with the COI amplicons.

### Sequence analysis

The initial read quality assessment was performed using FastQC v0.11.9 (Andrews, 2010) and summarized with MultiQC v1.13 (Ewels et al., 2016), which guided the selection of quality filtering parameters. Adapter and primer sequences were removed using Fastp v0.23.2 (Chen et al., 2018) (with the options -detect_adapter_for_pe, -disable_quality_filtering, and -disable_length_filtering) and Cutadapt v3.5 (Martin, 2011), applying linked adapters, discarding untrimmed reads, a minimum overlap of 10 bp, and a minimum read length of 1 bp. Subsequently, reads were truncated using the truncLen parameter (228 bp for R1 and 219 bp for R2 in 16S rRNA gene amplicons and 210 bp for R1 and 200 bp for R1 for COI gene amplicons), and filtering was applied based on maximum expected error thresholds of 2 for forward reads and 5 for reverse reads. Sequence denoising and chimera detection were carried out with DADA2 v1.22.0 (Callahan et al., 2016). Forward and reverse reads were merged using the DADA2 core sample inference algorithm with pooled inference to generate amplicon sequence variants (ASVs). Non-chimeric ASVs were taxonomically assigned using the IDTAXA classifier implemented in the DECIPHER v2.22 (Wright, 2016) package, applying a confidence threshold of 50%. Classification of the 16S rDNA amplicons was performed using the SBDI Sativa curated 16S rDNA GTDB reference database enriched with mitochondrial and chloroplast sequences and previous novel bacterial symbionts detected in nudibranch in our previous studies: *Candidatus* Doriopsillibacter (Džunková et al., 2023), *Ca*. Felimaribacterales (NCBI Assembly: GCA_977061665), *Ca*. Polyceribacterales (NCBI Assembly: GCA_977061625), *Ca*. Acroporibacterales (BioSample: SAMN55002297) (Porcel Sanchis et al., 2026). The COI amplicons were classified using the Metazoa COI Database (Heller et al., 2018). BLASTn analyses were also performed against the NCBI nt database in September 2026. Species-level assignments were accepted only when the best hits showed high sequence identity and consistent taxonomic assignment among the closest matches (Pappalardo et al., 2025). In COI amplicon sequencing output, reads assigned to the host *G. atlanticus* and species considered laboratory contamination (mice and *Polycera, Felimare, Peltodoris* nudibranch species processed routinely in our laboratory) were removed.

Core microbiomes of *P. porpita* and *G. atlanticus* were analyzed using the R package Microbiome v1.18.0 (Lahti and Shetty, 2017). For *G. atlanticus*, the core microbiome analyses were performed at the sample type level. Core ASVs were defined as those with a relative abundance >1% present in more than 50% of samples within each group. Principal component analysis (PCA) was conducted after filtering out ASVs with a relative abundance <0.01% and applying centered log-ratio (CLR) transformation to assess differences between sample groups. Core ASVs that could not be classified by IDTAXA classifier, were further examined using BLASTn against the NCBI database (accessed March 2026) (Altschul et al., 1990).

## Results and Discussion

### Alternative prey species detected

COI metabarcoding resulted in 6,927.89 ± 3,605.13 reads per sample (**Supplementary Table S1**), yielding 80 ASVs, of which the vast majority (98.93 ± 2.22%) were assigned to *G. atlanticus* (**Supplementary Table S2**). Despite the high amount of *G. atlanticus* DNA in its gut samples, two cnidarians were detected: *Pelagia noctiluca* (order *Semaeostomeae*, class *Scyphozoa*; average relative abundance of 0.07% in 1 individuals) and *Porpita porpita* (order *Anthoathecata*, class *Hydrozoa*; 0.038 and 0.050%, 2 individuals) (**Figure 2a**). Thus, despite the strong dominance of host DNA, our COI metabarcoding approach enabled the detection of prey-derived sequences, consistent with previous studies using host-blocking primers in fish, reptiles, birds, and mammals (Clark et al., 2023; Coveney et al., 2025). The near absence of detectable prey DNA in our samples is unlikely to be explained solely by methodological limitations, as *P. physalis* DNA was successfully detected at different low levels of dilution (1:100, 1:1,000, 1:10,000, 1:100,000, and 1:1,000,000) in the presence of *G. atlanticus* DNA (**Figure 2b**). In particular, the very low level of prey DNA in gut samples may indicate that the nudibranchs had not fed recently, possibly due to long transport to the beach, resulting in food deprivation for hours, days, or even weeks. Nevertheless, in the absence of prey, cannibalism, which has been described in some nudibranch species (Macali et al., 2023; Megina et al., 2002), cannot be ruled out, as its occurrence in *G. atlanticus* remains unknown.

**Figure 2.**
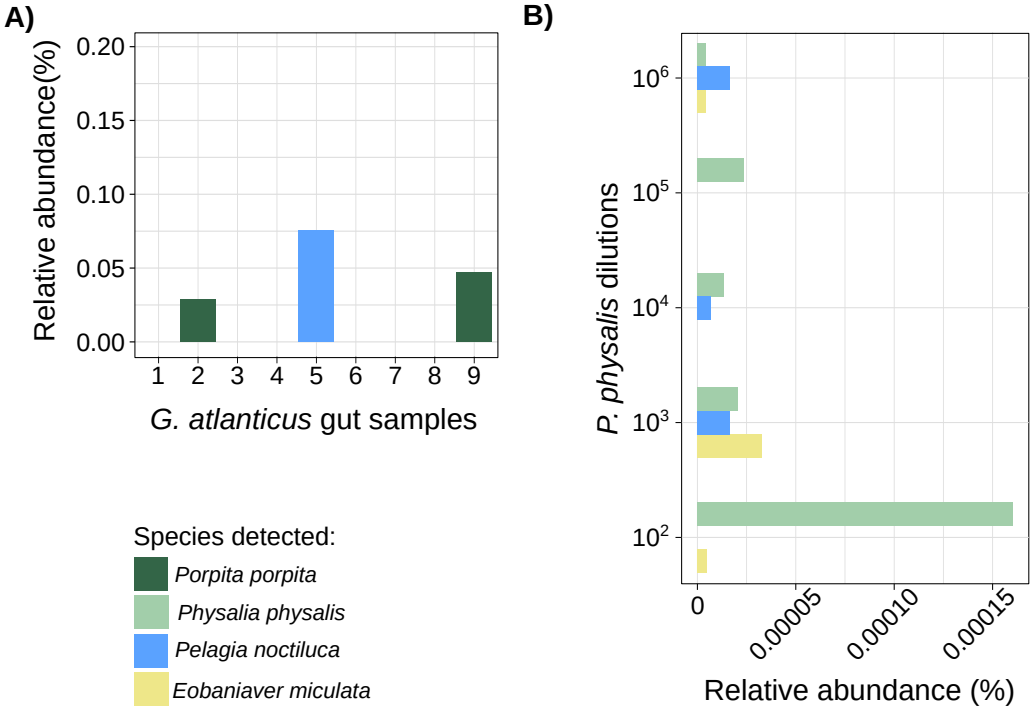
Analysis of prey-derived DNA in *G. atlanticus* samples. **a)** COI metabarcoding results for cerata tips and digestive tract samples. *P. noctiluca* was detected in two samples, while *P. noctiluca* in one sample. **b)** Relative abundance (%) of the non-*G*.*atlanticus* species in the COI metabarcoding experiment, in which *G. atlanticus* DNA was mixed with *P. physalis* DNA at 100-, 1,000-, 10,000-, 100,000-, and 1,000,000-fold dilutions. This positive control experiment demonstrates the ability of the assay to detect *P. physalis* DNA when its proportion is as low as 1,000,000-fold lower than that of *G. atlanticus* DNA. Note: *P. noctiluca* was naturally present in the *G. atlanticus* gut sample used for this assay, while *Eobania vermiculata* was associated with the *P. physalis* samples.

Our findings suggest that *G. atlanticus*, in the absence of its primary prey *P. physalis* in this area of Mediterranean, has broadened its trophic spectrum to include other available marine taxa, confirming the previously reported *P. porpita* as an alternative prey, and identifying an additional new prey species: *P. noctiluca*. This prey switch may reduce potential public health risks, as no adverse health effects have been reported following contact with the hydrozoan *P. porpita*, and the common Mediterranean jellyfish *P. noctiluca* is not considered highly hazardous, thus its presence does not lead to beach closures (Ballesteros et al., 2022; Hall et al., 2018).

Although we did not detect any traces of *P. physalis* DNA in *G. atlanticus* samples, more research is needed to evaluate the persistence of *P. physalis* nematocysts in *G. atlanticus*. Nudibranch kleptocnides can remain sequestered within their tissues for periods ranging from several days to weeks (Greenwood, 2009). Even if *G. atlanticus* appears in the Mediterranean at locations where no *P. physalis* is present, it may still carry nematocysts acquired from prey consumed in the Atlantic Ocean.

### G. *atlanticus* does not acquire microbes from *P. porpita*

Microbiome samples from *G. atlanticus* and *P. porpita* yielded a total of 2,123 - 46,956 reads per sample (average 13,841.10) (**Supplementary Table S1**), resulting in 1,049 ASVs (**Supplementary Table S3**), of which 685 were unique to *G. atlanticus* and 111 unique to *P. porpita* (**Figure 3a**). Each *G. atlanticus* body part had 63 - 163 specific ASVs. Only 62 ASVs were shared between the four different *G. atlanticus* body parts and *P. porpita*. The most abundant phylum across all *G. atlanticus* body parts and *P. porpita* was *Pseudomonadota* (27.01, 1.41-92.07% across all samples), followed by *Bacillota* (3.28, 0-27.29%) and *Actinomycetota* (2.08, 0-14.61%), which is a microbial composition commonly reported for other marine invertebrates (Ochoa-Sánchez et al., 2023) (**Supplementary Figure S2**).

**Figure 3.**
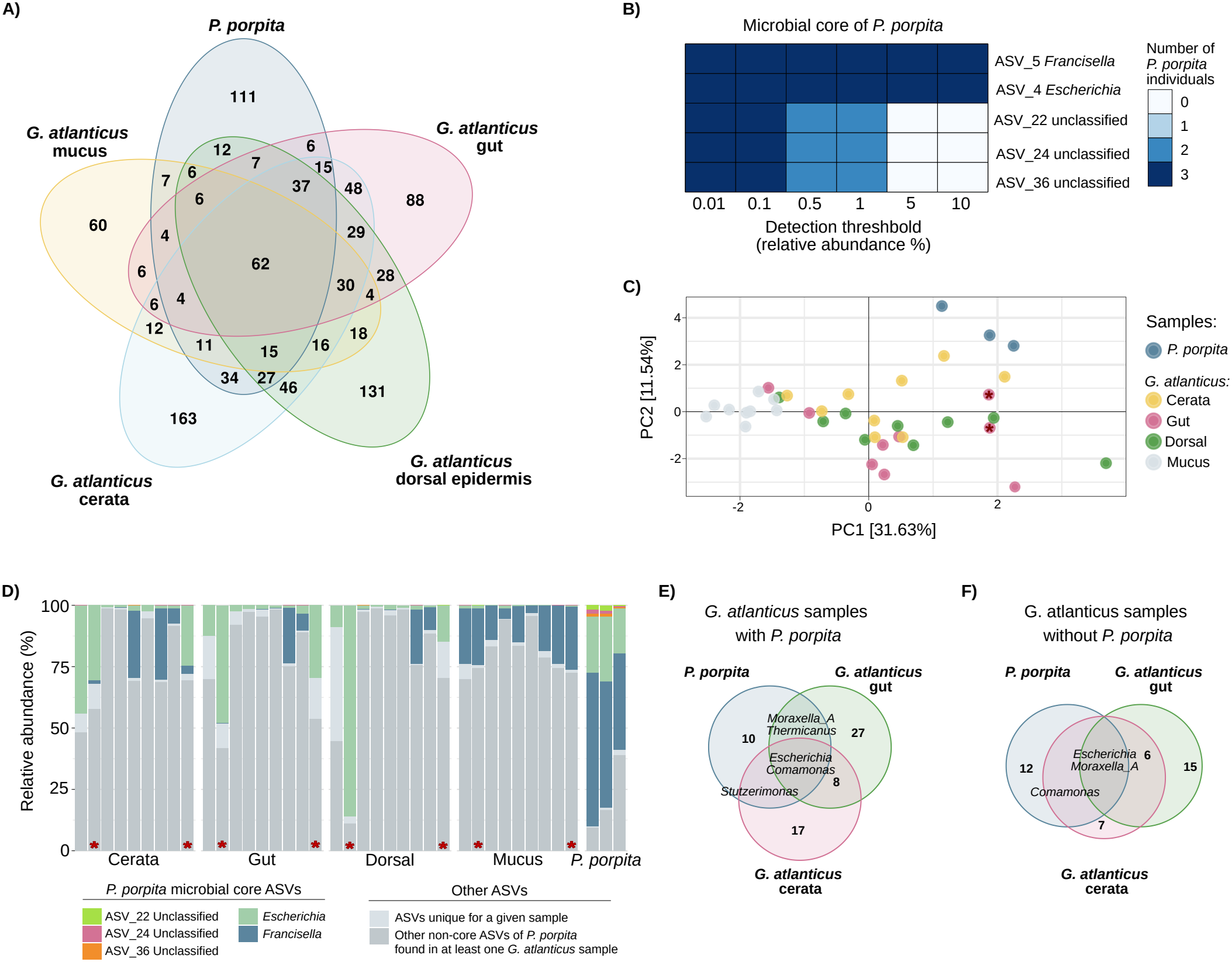
Microbiome analysis of *G. atlanticus* and *P. porpita*. **a)** Venn diagram showing the number of microbial ASVs detected across different *G. atlanticus* body parts and *P. porpita*. **b)** Core microbiome of *P. porpita*, showing the number of individuals in which core ASVs were detected under different relative abundance thresholds. **c)** Principal component analysis (PCA) of *P. porpita* and different body parts of *G. atlanticus*. Samples in which *P. porpita* traces were detected are marked with a star. **d)** Proportion of core *P. porpita* ASVs detected in *G. atlanticus* individuals in which *P. porpita* traces were identified by COI metabarcoding. The barplots also show proportion of ASVs, which do not belong to *P. porpita* core microbiome, but were found in some *G. atlanticus* individuals. Additionally, ASVs unique to each sample are shown. **e)** Venn diagram of microbial ASVs with relative abundance >1% in *P. porpita* samples and in samples of gut, cerata and mucus of *G. atlanticus*containing traces of *P. porpita* DNA . **f)** Venn diagram of microbial ASVs with relative abundance >1% in *P. porpita* samples and in samples of gut, cerata and mucus of *G. atlanticus* individuals without detected *P. porpita* DNA.

In *P. porpita*, among the taxa with the average relative abundance >1% were *Escherichia* (20.4 ± 4.87%), *Francisella* (50.26 ± 11.59%), *Thermicanus* (average 2.15, range 0-6.4%), *Acinetobacter* (1.90, 0-4.40%), *Anoxybacillus* (1.48, 0-4.45%), and unclassified ASV_22, ASV_24 and ASV_36 (**Supplementary Table S4**). However, among these taxa, only *Escherichia, Francisella* and unclassified ASV_22, ASV_24 and ASV_36 were part of the *P. porpita* core microbiome at the 1% threshold, suggesting that most of the taxa detected in *P*.*porpita* represent transient or non-specific microbial associations (**Figure 3b**). Consistent with this pattern, *P. porpita* exhibited a higher Shannon diversity index than *G. atlanticus* (median 2.25 vs. 1.54), reflecting an even distribution of numerous low-abundance taxa rather than dominance by a few lineages, with values falling within the range reported for other hydrozoans such as *Hydra, Millepora*, and *Lytocarpia* (McCauley et al., 2023). These patterns likely reflect the ecology of hydrozoans, as the surface-dwelling pleustonic lifestyle of *P. porpita* increases exposure to diverse environmental microbes, including through contact with beach sand during stranding.

The microbial composition of *G. atlanticus* samples did not include a substantial proportion of the *P. porpita* core microbiome (**Figure 3c, 3d**), suggesting limited microbial transfer. In the PCA (**Figure 3c, Supplementary Figure S3**), only three cerata samples and one digestive tract sample clustered within the region occupied by *P. porpita* samples, whereas mucus samples were positioned furthest from *P. porpita* in ordination space. This pattern suggests that, if any bacteria are transferred from prey, they are not retained in both the mucus and the cerata, as was observed in previous studies (Zhukova et al., 2022; Džunková et al., 2023). *Escherichia* and *Francisella* were the most abundant bacterial taxa in *P. porpita* and were also detected in several, but not all, *G. atlanticus* samples (**Figure 3d**). It is unlikely that *Escherichia* and *Francisella* are symbionts involved in chemical defense transferred from *P. porpita* to *G. atlanticus*, given that members of these genera include opportunistic or pathogenic bacteria associated with a range of animal hosts, have been detected in association with reduced coral health, and can persist in diverse environmental habitats (Gignoux-Wolfsohn et al., 2017; Jang et al., 2017; van Elsas et al., 2011). Moreover, the microbial composition of the dorsal central body, which served as a negative body-part control because symbionts acquired through nematocysts would not be expected to occur there, was similar to that of the cerata (**Figure 3c, 3d**), which would not be expected to contain bacteria involved in nudibranch chemical defense.

As additional evidence against the transfer of mutualistic symbionts involved in chemical defense, the microbiome of all nudibranch body parts did not differ between individuals in which *P. porpita* DNA was detected and those in which no traces of *P. porpita* were found (**Figure 3d, 3e, 3f)**. In individuals in which *P. porpita* DNA was detected in the gut, *Escherichia* (average relative abundance 22.88%, range 1.3–48.09%), *Comamonas* (3.53%, 0.06–9.55%), *Moraxella* (1.19%, 0.02–4.68%), *Thermicanus* (0.22%, 0–1.08%), and *Stutzerimonas* (0.5%, 0-2.098%), were the only taxa with relative abundance of >1% found to be shared with *P. porpita* (**Figure 3e**). The same taxa were also detected in the individuals in which no *P. porpita* DNA was detected (**Figure 3f**). Although mutualistic bacteria could theoretically have stably colonized *G. atlanticus* individuals, making them detectable despite the absence of prey in the gut at the time of sampling, the detected bacterial taxa exhibit broad environmental distributions and are frequently reported in association with opportunistic pathogenicity in mammals, molluscs, and fish (An et al., 2008; Goldstein et al., 2009, 2009; Ryan et al., 2022). Overall, these results provide no clear evidence that the shared bacterial taxa were transferred from *P. porpita* to *G. atlanticus*, although their environmental origin could have been further investigated by comparing the microbial communities of beach sand or seawater collected at the same location.

As none of the microbial taxa shared between *P. porpita* and *G. atlanticus* belonged to lineages typically associated with stable mutualistic interactions (**Figure 3**), we next focused on identifying potentially beneficial core microbes across different body parts of *G. atlanticus* that may be established independently of its prey (**Figure 4**). Microorganisms contributing to host defense would be expected to form core microbiome of the exposed tissues, such as mucus and cerata, which are directly involved in predator interactions, and comparatively depleted in the central dorsal body surface, used here as a negative control. Although each of the four sampled body parts harbored distinct microbial taxa (**Figure 3a**), most of the detected taxa were present at low relative abundances, whereas symbionts involved in chemical defense are typically expected to be both abundant and consistently detected across individuals. Therefore, we focused on core microbiome analyses for each body part, defined as bacteria with a relative abundance >1% present in more than 50% of samples.

**Figure 4.**
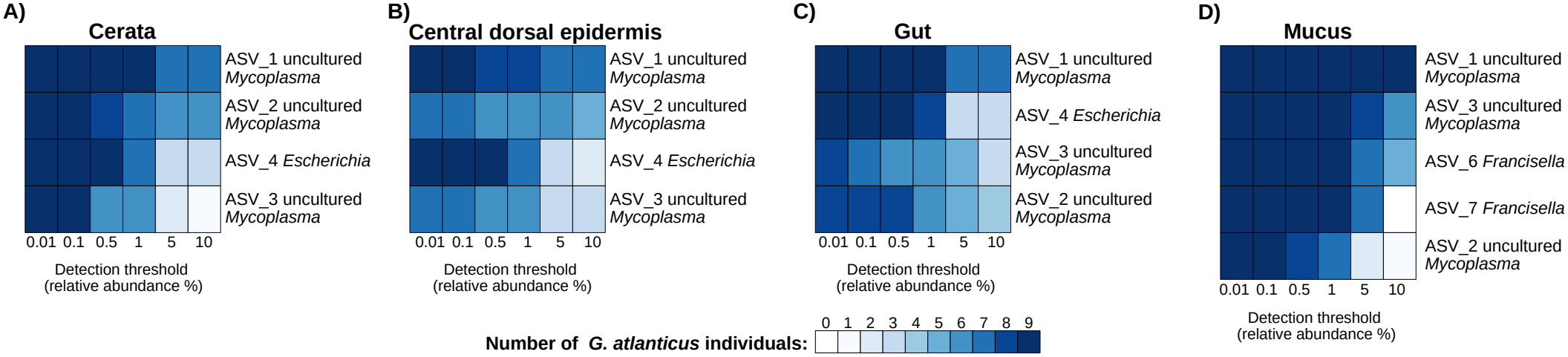
Core microbiomes of *G. atlanticus* body parts. Diagrams show **a)** cerata, **b)** central dorsal epidermis, **c)** gut, and **d)** mucus. Each diagram indicates the number of individuals in which core ASVs were detected under different relative abundance thresholds.

The core microbiome shared across all *G. atlanticus* samples comprised three ASVs assigned to uncultured *Mycoplasma* (*Mollicutes*) (64.07 ± 32.80%, cumulative relative abundance of ASV_1, ASV_2 and ASV_3), *Escherichia* (ASV_4 average abundance 9.23%, range 0.11-85.91%), and two *Francisella* ASVs (7.76 % cumulative average abundance of ASV_6 and ASV_7, range 0-28.90%) (**Supplementary Table S4, Supplementary Figure S4**). The highest abundance of *Mycoplasma* and *Francisella* was detected in mucus (75.82 ± 11.17% and 16.65 ± 8.36%, respectively), while the highest abundance of *Escherichia* was detected in cerata and gut (44.0-0.74% and 48.08-0.95%).

When core microbiomes were defined separately for each of the four body parts, *Mycoplasma* and *Escherichia* were consistently identified in the cerata, central body, and gut (**Figure 4a-c**), while the mucus core microbiome additionally included *Francisella* (**Figure 4d**). *Mycoplasma, Escherichia* and *Francisella* are unlikely to represent mutualistic symbionts involved in host defense, given their broad environmental distribution and the presence of species known to be pathogenic to humans (Degabriel et al., 2023; Lanao et al., 2026; Naidoo and Zishiri, 2025), marine mammals (Duarte-Benvenuto et al., 2023), mussels (Grkovic et al., 2021), and fish (Rocha et al., 2025). Furthermore, mutualistic bacteria involved in the chemical defense of the nudibranch are expected to form part of the core microbiome of the cerata and mucus, and could also be detected in the gut, but not in the central dorsal body, which does not store nematocysts. Since the four body parts contained nearly the same core microbiome, we found no evidence of a body-part-specific bacterial community that could be associated with chemical defense. While our study, based on previous knowledge of defense systems in dorid nudibranchs, focused on identifying potential microbial mutualists that may produce defensive molecules for their aeolid nudibranch host, further research should also explore microbes that may produce precursors of these defensive molecules.

Collectively, these results indicate that the dominant and core microbial taxa associated with *G. atlanticus* are unlikely to play a central role in host chemical defense, reinforcing the absence of evidence for functionally relevant mutualistic symbionts in this system.

## Conclusions

This study provides the first molecular characterization of the diet of *G. atlanticus*, with the aim of assessing also whether microbial symbionts involved in chemical defense may be acquired from prey. While *G. atlanticus* typically derives its defensive capacity from nematocysts obtained from the venomous *P. physalis*, our results indicate that, in the absence of this primary prey, the nudibranchs shift to alternative cnidarians *P. porpita* and *Pelagia noctiluca* commonly found along the Mediterranean coasts that are not considered highly hazardous.

Microbiome analyses further revealed no evidence for the transfer of potentially mutualistic bacteria from *P. porpita* to *G. atlanticus*. Instead, the microbial communities associated with different body parts were dominated by taxa with broad environmental distributions, including lineages frequently reported as opportunistic pathogens. These findings indicate that, in contrast to some dorid nudibranchs, the aeolid *G. atlanticus* does not appear to rely on microbial symbionts for the production of defensive compounds.

## Data availability

All data have been submitted to EMBL-EBI within the Bioproject PRJEB112072 and PRJEB128141.

## Acknowledgments

We would like to thank the Seqplexing S.L., Valencia, for DNA sequencing. The computations were performed on the HPC cluster Garnatxa at the Institute for Integrative Systems Biology (I2SysBio), which is a mixed research center formed by the University of Valencia (UV) and Spanish National Research Council (CSIC).

## Authors Contribution

Wrote the manuscript: DPS, MD. Provided samples: SP, IAT, LP. Extracted DNA and prepared amplicons: AOM, IAM, MD. Analyzed DNA sequence data: DPS. Performed additional analysis: CA. Provided funding and supervised the study: MD.

