## Supplementary Figures for "The blue dragon nudibranch (*Glaucus atlanticus*) shifts to less venomous prey in the Mediterranean, with no evidence of defensive microbial symbionts"

### The blue dragon nudibranch (*Glaucus atlanticus*) shifts to less toxic prey in the Mediterranean and shows no microbial contribution to chemical defense

Dafne Porcel Sanchis, Alicia Ortiz-Maiques, Ines Alberola Mora, María Asunción Piedecausa Narejos, Conxita Avila, Mária Džunková

**Supplementary Figure S1**

Agarose gel showing PCR amplification with universal COI primers compared to reactions including blocking primers. Samples included G. atlanticus (S), aphid DNA (control C+), and a no-template control (C−). Universal COI primers produced amplification in both animal DNA samples (S and C+). In reactions containing a 5:1 ratio of blocking to universal primers, amplification of G. atlanticus DNA was strongly reduced, while aphid DNA remained detectable, although with decreased amplification due to the lower concentration of universal primers. MW - molecular weight marker (in bp).

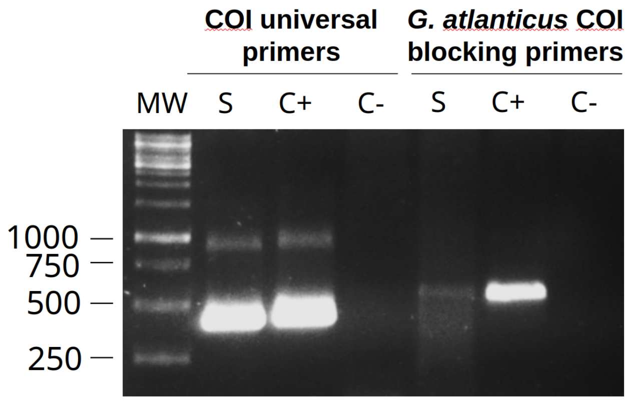

**Supplementary Figure S2**

Relative abundances of ASVs on phylum and genus level in *G. atlanticus* samples (mucus, central dorsal epidermis, gut, cerrata) and in *P. porpita.*

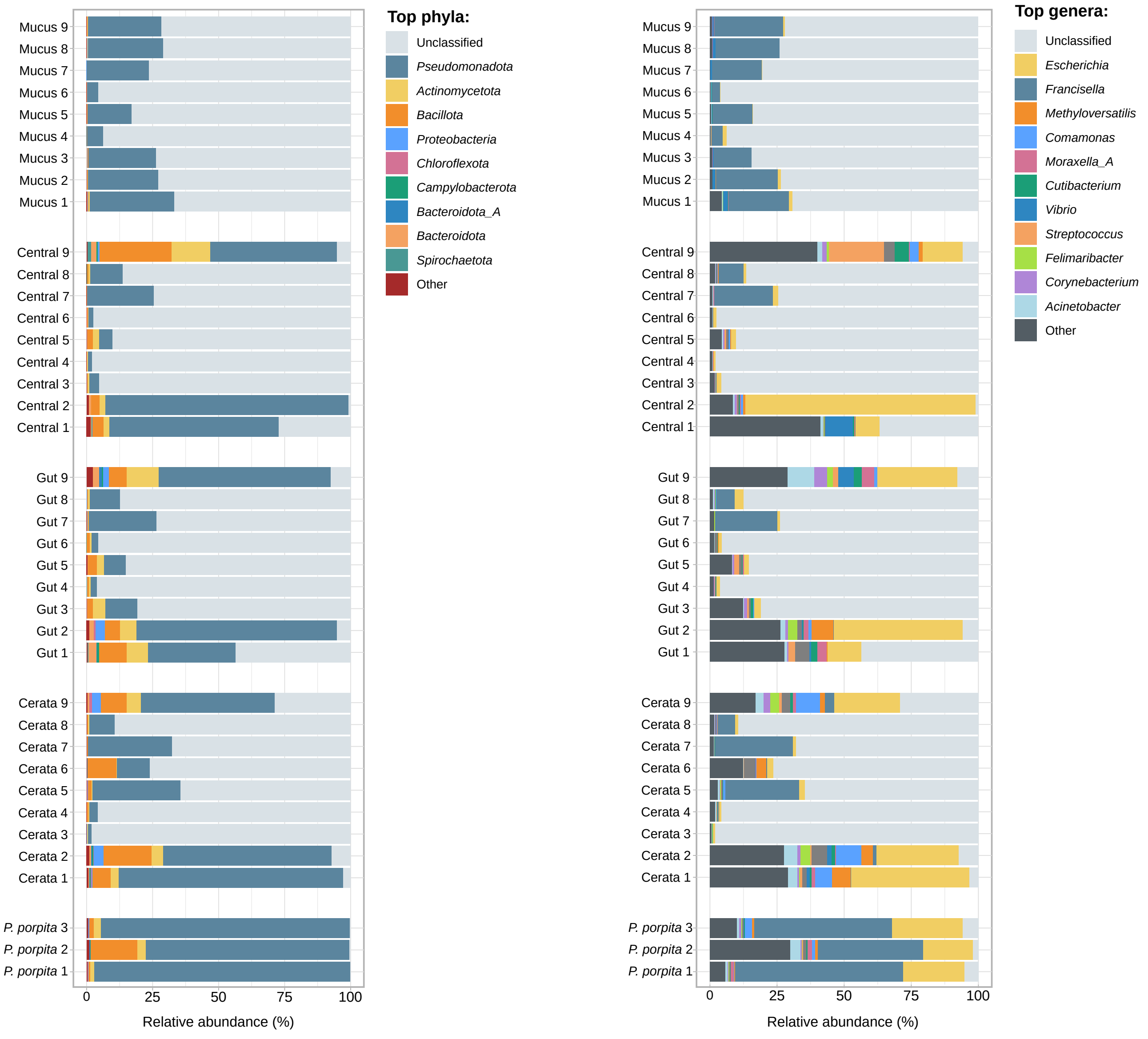

**Supplementary Figure S3**

Bacterial ASVs contributing to sample ordination along PCA axes. The figure shows the ASVs with the highest loadings on PC1 and PC2, colored on family level.

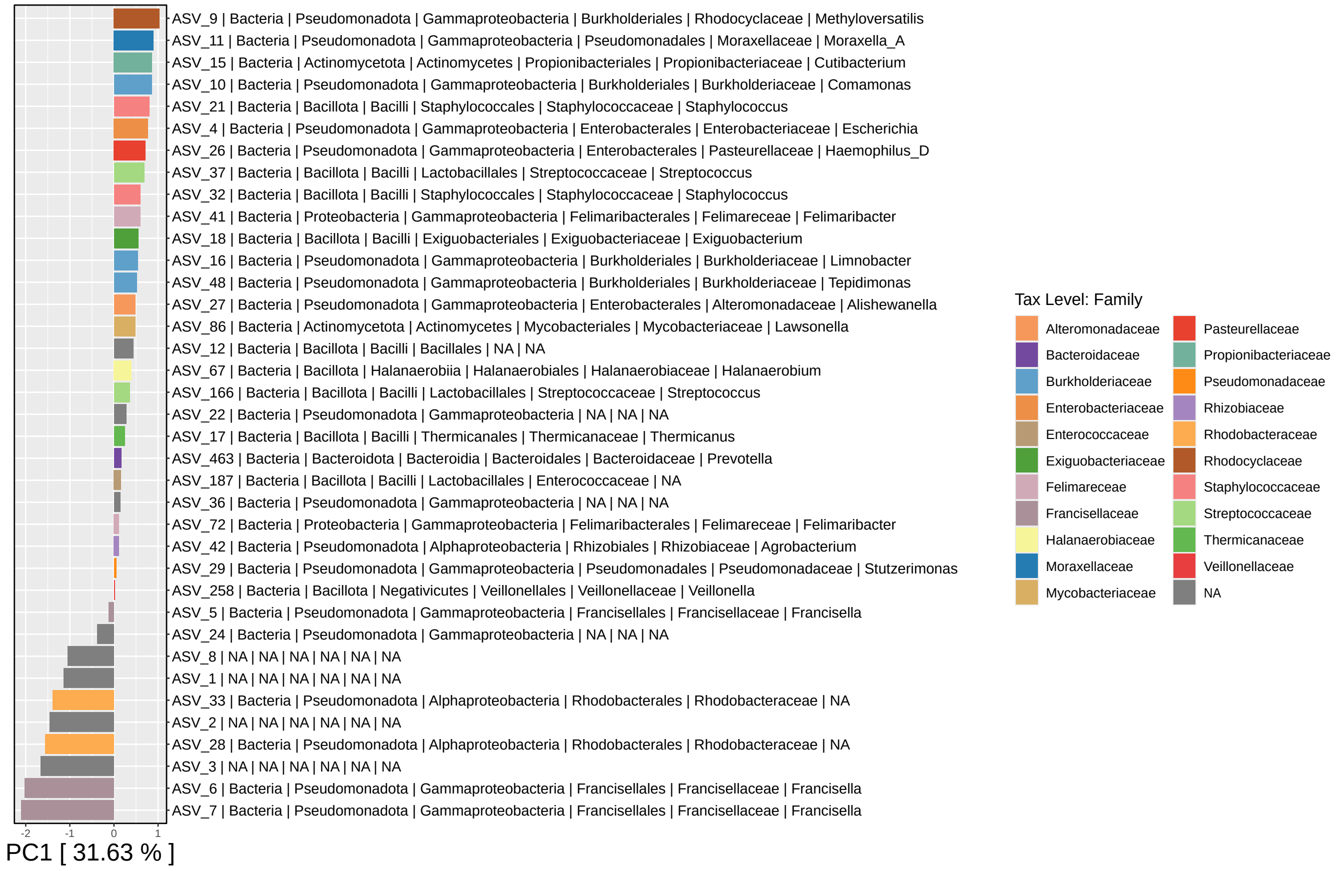


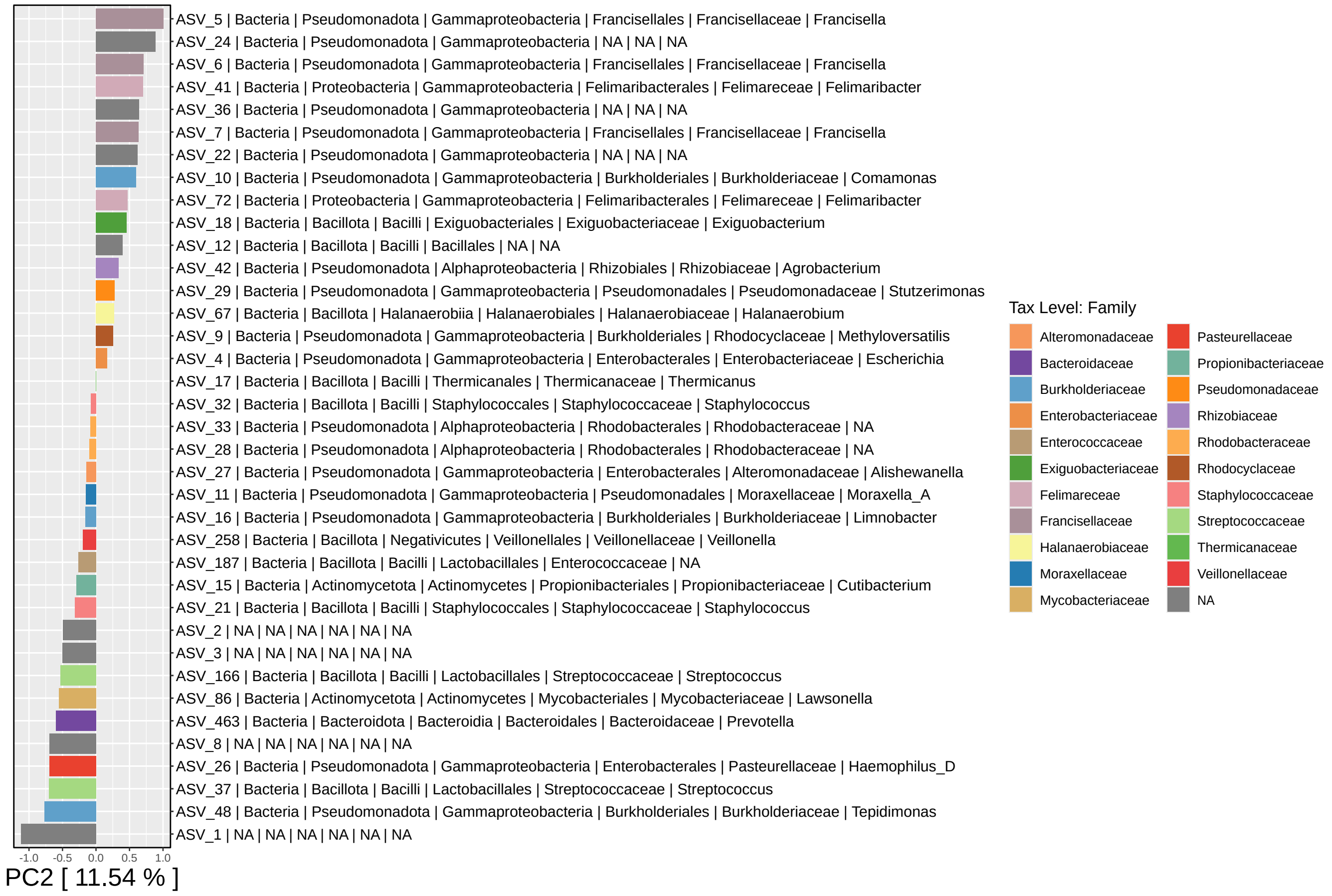

**Supplementary Figure S4**

Core microbiome of all *G. atlanticus* samples. The diagram indicates the number of samples, in which core ASVs were detected under different relative abundance thresholds.

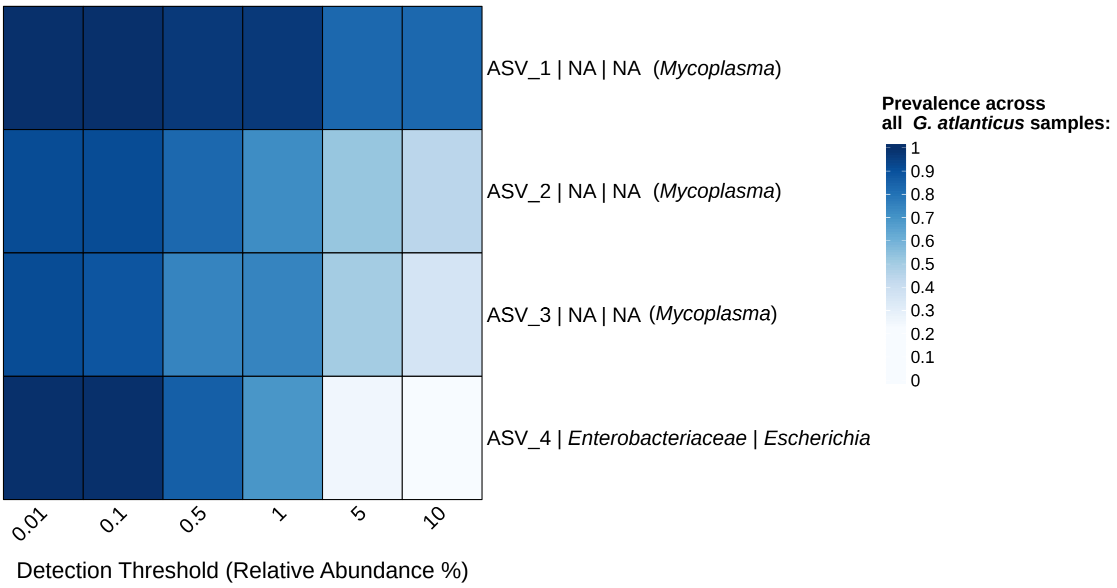
